# Prot2Token: A multi-task framework for protein language processing using autoregressive language modeling

**DOI:** 10.1101/2024.05.31.596915

**Authors:** Mahdi Pourmirzaei, Farzaneh Esmaili, Mohammadreza Pourmirzaei, Duolin Wang, Dong Xu

## Abstract

This paper proposes a versatile tokenization method and introduces Prot2Token, a model that combines autoregressive language modeling with protein language models (PLMs) to tackle various protein prediction tasks using protein sequences. Leveraging our tokenization method, Prot2Token adapts existing PLMs for multiple tasks such as protein-level prediction, residue-level prediction, and protein-protein interaction prediction through next-token prediction of tokenized target label sequences. By incorporating prompt tokens into the decoder, Prot2Token enables multi-task training in a single end-to-end session. Our results demonstrate that Prot2Token not only matches the performance of specialized models across various tasks but also paves the way for integrating protein tasks with large language models (LLMs), representing an important step towards creating general-purpose PLMs for advanced protein language processing (PLP). Additionally, we use Prot2Token to develop S-ESM, a structure-aware version of the ESM model, which achieves competitive performance with state-of-the-art methods in 3D structure-related tasks using only protein sequences. Code is available at: https://github.com/mahdip72/prot2token.

## 1. Introduction

Proteins, with their vast diversities and functions, are fundamental to biological research and medicine; yet our understanding of them remains incomplete. A crucial aspect of this understanding is the deep learning representation of protein sequence, which aids in predicting protein functions, identifying protein-protein interactions, and designing novel proteins (Shim et al., 2019; Manshour et al., 2023). Building upon this, protein language models (PLMs) have emerged as powerful tools in protein language processing (PLP), which applies language modeling and other natural language processing (NLP) techniques to decipher the language of amino acid sequences in terms of protein properties and behavior (An & Weng, 2022). This capability has resulted in their superior performance across various tasks related to protein functions and interactions prediction (Rives et al., 2021; Elnaggar et al., 2021).

Despite the impressive capabilities of PLMs in various protein-related tasks, there is still a lack of a unified framework that can effectively address the diverse range of advanced PLP tasks. Existing PLMs are often developed to be task-specific, requiring separate architecture design and training for each task, which can be time-consuming and computationally expensive. In addition, they can not handle different types of PLP tasks at the same time (Hsu et al., 2022; Hu et al., 2023; Roche et al., 2023), a crucial aspect of creating a general-purpose PLM.

After the success of autoregressive large language models (LLMs), there has been tremendous work to utilize LLMs beyond NLP, to other modalities (Kondratyuk et al., 2023; Lu et al., 2023; El-Nouby et al., 2024; Liu et al., 2023). One thing that these methods all have in common is treating all the targets as sequences and using the simple next token prediction loss function to train. This makes the labels from different tasks to be encoded into a unified sequence of tokens. In other words, every label that can be encoded into fixed-sized tokens can be handled by a unified LLM. Drawing from this inspiration, we propose a unified strategy for tokenization and introduce Prot2Token, which merges pretrained PLMs with an autoregressive language modeling decoder to do PLP. Prot2Token can be connected to existing PLMs and align them to predict different types of tasks given protein sequences, as shown in Figure 1. By adopting next-token prediction as the learning objective and using task prompts for the guidance of prediction, this method can also harness the strengths of multi-task representation learning to enhance performance and generalization while reducing the need for labeled training data across various protein prediction tasks (Vandenhende et al., 2021).

**Figure 1.**
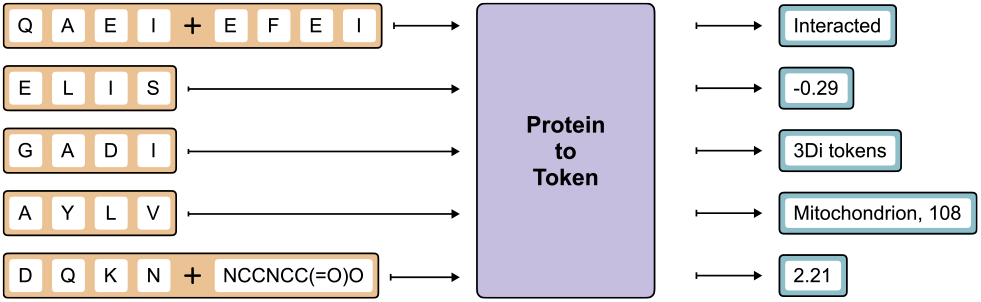
Overview of the Prot2Token process. The model accepts protein and SMILES sequences as the input and predicts corresponding labels across various PLP tasks.

In this paper, we showed that Prot2Token can be built on top of ESM-2 (Lin et al., 2023) models to be a substitute for the current highly specialized models with similar performance. It can be used as either one-task learning or jointly trained with multiple tasks in an end-to-end fashion. Additionally, our findings suggest that Prot2Token’s effectiveness can be increased when addressing tasks with limited data samples by integrating auxiliary tasks—either related supervised tasks or synthetic self-supervised ones alongside the main task.

Prot2Token is not limited to prediction and can be used for other purposes such as aligning existing PLMs to be structure-aware via training on 3D structure tokens as the label, inspired from a series of recent works (Heinzinger et al., 2023; Su et al., 2023). That is, we extended our work by making ESM to be a structure-aware ESM, named S-ESM, via predicting FoldSeek (van Kempen et al., 2023) 3Di tokens given protein sequences and demonstrate that despite the simplicity, it can significantly outperform the original ESM on 3D structure-related PLP tasks. Prot2Token is a step towards aligning autoregressive models for advanced PLP and building general dialogue-based protein language models.

The contribution of this work can be summarized as: (1) We propose a novel tokenization strategy for advanced PLP tasks, including protein-level prediction, residue-level prediction, and protein-protein as well as protein-ligand interaction prediction, and design a model named Prot2Token. Prot2Token can be applied to existing pre-trained PLMs and align them with multiple PLP tasks in an end-to-end fashion. (2) We show that Prot2Token can effectively perform multi-task learning, demonstrating that predicting PLP tasks through Prot2Token often benefit from simultaneously learning multiple tasks. (3) Using Prot2Token, we upgrade the ESM-650m model to be structure-aware, named S-ESM, and show that it improves the ESM on 3D-informed protein tasks.

## 2. Related works

The methods related to PLMs can be broadly classified into three primary categories: sequence-based models, structure-based models, and models that integrate both sequence and structure information. PLMs are becoming increasingly significant and popular in biological research, particularly for tasks related to protein prediction (Lin et al., 2023; Elnaggar et al., 2021). In addition, a few studies have explored dialogue-based protein language models for PLP tasks and *de novo* protein generation (Lv et al., 2024; Wang et al., 2023). For a more detailed discussion of these categories and related work, refer to Appendix Section A.1.

## 3. Method

### 3.1. Tokenization

Prot2Token consists of two sections, two encoders, and one autoregressive decoder, meaning we have three tokenizers. As for the encoders, we utilize the original tokenizers from both pre-trained ESM-2 and BARTSmiles (Chilingaryan et al., 2022) models, along with their pre-trained embedding layers. For the details regarding the encoders tokenizer, refer to Appendix Section A.2.1. The rest of this section is about building a tokenizer for the autoregressive decoder part of Prot2Token. In the first step of the tokenization process of PLP labels, we incorporate two special tokens into the tokenizer: *<BOS>* refers to the beginning-of-sequence, and *<EOS>* refers to the end-of-sequence, into the tokenizer. These tokens are important for restricting the start and the finish of the output of the sequence by the decoder. Furthermore, the key step in the Prot2Token model is to convert every type of label into a sequence of discrete tokens. In the domain of protein studies, we encounter multiple types of tasks, each requiring customized treatment. To find the details of each one, refer to Appendix Section A.2.2. In the end, we convert all output tokens (labels) as well as task tokens (Section 3.2) to trainable embedding vectors before passing them into the decoder.

### 3.2. Architecture

The core idea of Prot2Token is to integrate an autoregressive decoder language model with existing encoder-style protein and chemical language models through cross-attention layers. This approach reformulates the labels of all tasks as sequences of tokens. In this framework, protein and chemical sequences are first processed by their respective encoders, transforming them into feature representations. These features are then fed into a decoder transformer, which predicts the labels sequentially. In the design of our model, inspired by Pix2Seqv2 (Chen et al., 2022), we introduce a mechanism to enhance the task adaptability of Prot2Token to solve multi-task at once. This is achieved through the introduction of a “task token” (prompt) at the beginning of each sequence label in the decoder part of the model. For more details about the architecture, refer to Appendix Section A.3. This strategy enables us to utilize a single decoder to predict outputs for each task, as demonstrated in Figure 2, thereby simplifying the inference process and reducing deployment cost. Mathematically, during the training process of the label sequence, while the task token is integral in directing the model, it is treated distinctly in terms of its weight assignment during loss calculation similar to Pix2Seq V2 method. Specifically, when calculating the likelihood of the protein sequence, we assign a zero weight to the prompt (task) token. Technical details of the weighting assignment are described in A.3.

**Figure 2.**
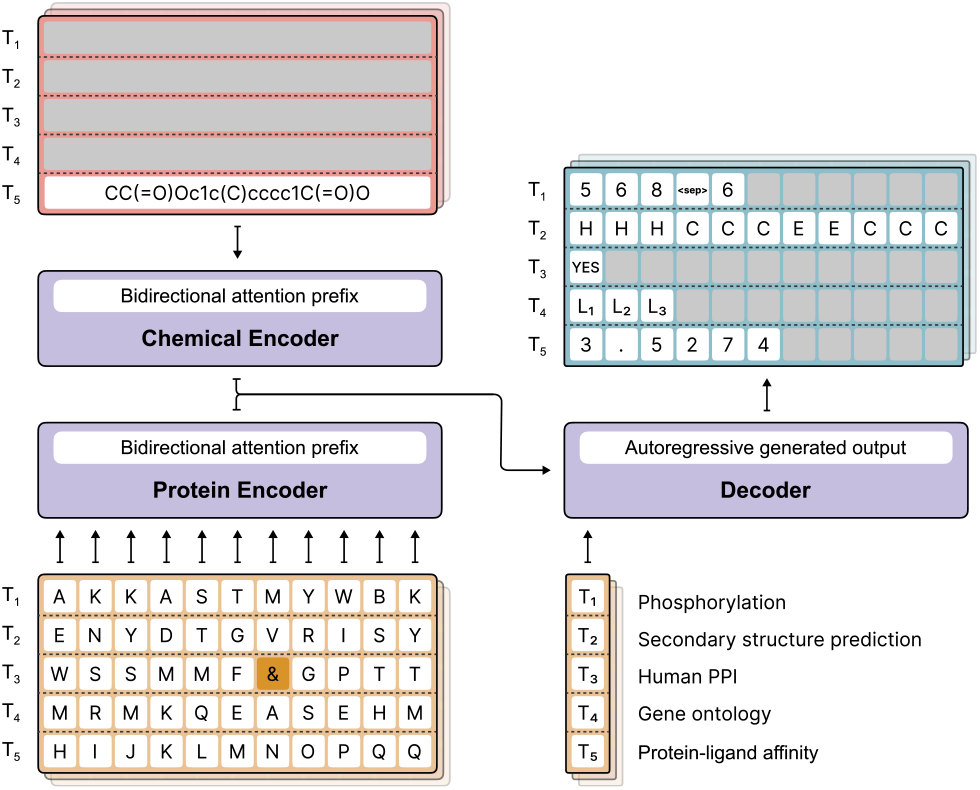
Training and prediction of multiple tasks using Prot2Token framework. This illustration demonstrates the capability of the Prot2Token model to be used concurrently on a variety of PLP tasks in a single end-to-end training. The encoders, with a bidirectional attention prefix, take in protein and chemical sequences and pass the encoded features to a transformer decoder. The decoder then generates output through an autoregressive process by conditioning on the task tokens. Each task token (T1 through T5) corresponds to a different task in the illustrated batch.

### 3.3. Datasets

In this work we consider several type of tasks and datasets from the benchmark of PEER (Xu et al., 2022), Protein-Shake (Kucera et al., 2023), CATH (Wang et al., 2024) and AlphafoldDB (Varadi et al., 2022), as well as other individual datasets. These datasets represent different types of tasks including regression, multi-class and multi-label classification, residue-wise classification, and sequence prediction. The detail of each dataset is placed in Appendix Section A.4.

## 4. Experiments

In this section, we initially demonstrated the application of Prot2Token for various downstream tasks, treating each type of task separately. Lastly, with the help of Prot2Token, we adapted the ESM model to be structure-aware, S-ESM, enhancing its capability to understand and utilize the 3D structure of proteins from sequences.

For all of our experiments, we considered ESM-2 (Lin et al., 2023) family of models as the protein encoder of Prot2Token. For the decoder part, we used an autoregressive language model with different configurations based on the size of the ESM encoder and hyperparameters of the autoregressive decoder (Appendix Section A.3). We only considered BARTSmiles as the chemical encoder for the protein-ligand affinity task and disabled it for the other tasks.

For all our experiments, we employed the Adam optimizer (Kingma & Ba, 2014) with a modification to decouple weight decay (Loshchilov & Hutter, 2017), setting beta-1 to 0.9 and beta-2 to 0.999. Our learning rate strategy was based on cosine annealing with initial warm-up steps (Loshchilov & Hutter, 2016). This approach was applied in all tasks. Additionally, all experimental protocols and models were developed using the PyTorch framework (Paszke et al., 2019).

### 4.1 Regression

This category includes three tasks: stability prediction, fluorescence prediction, and protein-ligand affinity prediction. The input is a protein sequence in the first two tasks, and the label is a floating-point number. For the protein-ligand affinity prediction, the input consists of both protein and molecule (SMILES) sequences, with the output being a floating-point number. The results are shown in Tables 1, 2, and 3. Additional details are in Appendix Section A.5. We beat the PEER methods in these predictions. Also, the fluorescence results showed that the performance boosts up to 5.6 percent by using multi-task learning (Table 2).

**Table 1.** Comparing Prot2Token with other methods on stability prediction.

| METHOD | SPEARMAN | MODEL |
| --- | --- | --- |
| BASELINE | 0.7527 | ESM-650M |
| PEER (FINE-TUNED) | 0.75 | ESM-1B |
| PEER (FINE-TUNED) | 0.771 | PROTBERT |
| OUR | 0.7947 | PROT2TOKEN (ESM-650M) |

**Table 2.** Comparing fluorescence prediction methods w/ and w/o multi-task learning. PLA and ST stand for protein-ligand affinity and stability, respectively. We considered the fine-tuned methods of PEER as the comparison.

| METHOD | AUX-TASKS | SPEARMAN | MODEL |
| --- | --- | --- | --- |
| PEER | - | 0.679 | ESM-1B |
| PEER | - | 0.679 | PROTBERT |
| OUR | - | 0.7389 | PROT2TOKEN (ESM-650M) |
| OUR | PLA | 0.7766 | PROT2TOKEN (ESM-650M) |
| OUR | PLA+ST | 0.78 | PROT2TOKEN (ESM-650M) |

**Table 3.** Comparing protein-ligand affinity prediction methods on the test set.

| METHOD | SPEARMAN | MODEL |
| --- | --- | --- |
| PEER (FINE-TUNED) | 1.559 | ESM-1B |
| PEER (FINE-TUNED) | 1.562 | PROTBERT |
| OUR | 1.3887 | PROT2TOKEN (ESM-650M) |

### 4.2. Classification

This category includes multi-class, multi-label and hierarchical classification tasks: Deeploc 2.0, enzyme reaction (ER) and TargetP localization, enzyme commission (EC), three types of gene ontology (GO) tasks, human protein-protein interaction (Human PPI), and fold classification. The results are shown in Tables 4 and 5. In Deeploc 2 dataset, we significantly improved the performance compared to the original method, and also, the ER task result showed that the performance boosted 7.5 percent by using multi-task learning. We could not calculate the Fmax metric for the EC and GO tasks, so we only considered the accuracy and F1 scores to evaluate performance. Consequently, direct comparisons with other methods were not possible. Additionally, we found that training on the fold classification dataset without incorporating auxiliary tasks was unstable, preventing the model from learning and producing the labels correctly. Supplementary results and additional details are in Appendix Section A.5.

**Table 4.**
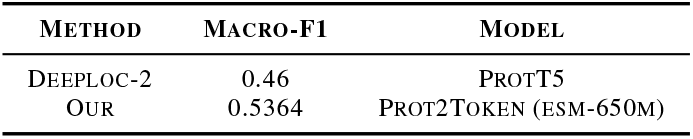
Localization prediction using Deeploc-2 dataset. The results are based on the independent test set.

**Table 5.**
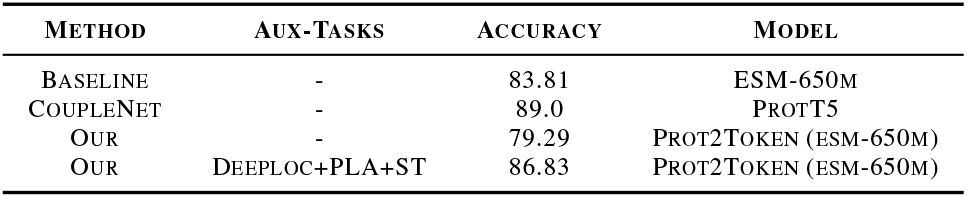
Comparing methods on ER dataset. PLA and ST stand for protein-ligand affinity and stability, respectively.

### 4.3. Sequence Prediction

This category includes different tasks from previous categories including secondary structure (SS) prediction, phosphorylation post-translational modification (PTM), Fold-Seek token-based 3D structure prediction, and protein-protein interface prediction. The result of SS (Table 6) showed competitive performance compared to the baseline. The results of additional tasks such as predicting FoldSeek tokens are in Appendix Section A.5.

**Table 6.**
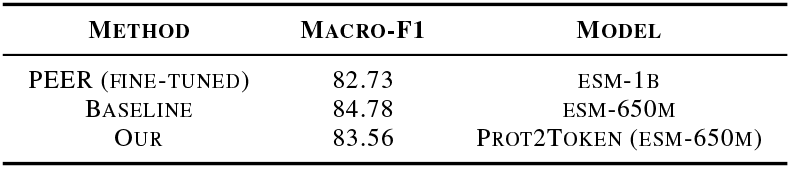
Secondary structure prediction evaluation. The baseline involves a linear classifier on top of the frozen ESM model.

### 4.4. Structure-Aware ESM

We found that Prot2Token excels in solving protein tasks and enhances the structure-awareness of current PLMs. While these models are typically trained on sequences alone, integrating structural information significantly improves performance on 3D-related tasks. Prot2Token could predict 3D structures by converting them into sequences of tokens, with FoldSeek being the most efficient method. We trained a model as described in Appendix A.6 and evaluated the fine-tuned encoder, S-ESM, using T-SNE. Figure 3 shows this evaluation. The results demonstrated that S-ESM could generate sequence embeddings aware of structural information. More details about this evaluation are in Appendix A.6.

**Figure 3.**
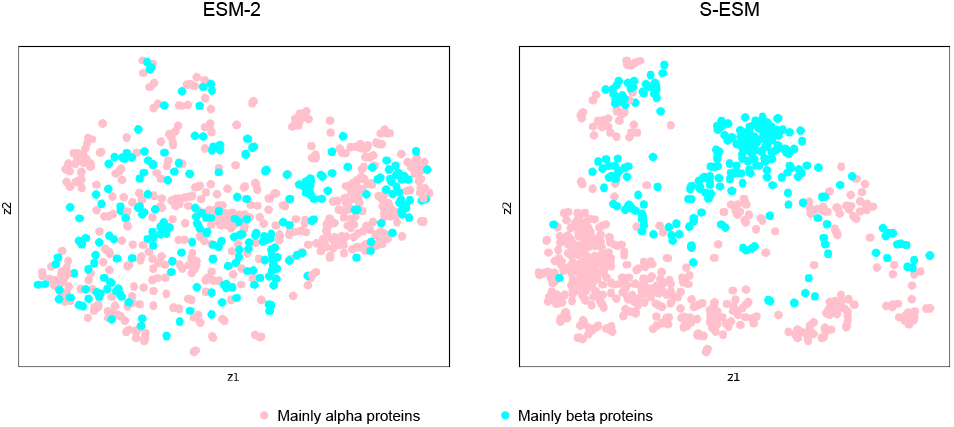
T-SNE visualization of sequence embeddings from ESM-2 and S-ESM for CATH structure domains. Both models are based on the ESM-650m architecture.

Additionally, we evaluated S-ESM on PLP tasks that require structural information. For this evaluation, we used the S-ESM (based on the ESM-650m architecture) as the encoder backbone. Following training on multiple tasks, we conducted a comparative analysis across various protein tasks, as detailed in Table 14 in Appendix Section A.5. The results demonstrated that S-ESM outperformed the original ESM in most cases and was competitive with state-of-the-art methods.

## 5. Discussion

Our study demonstrates the generalization of our tokenization framework in the Prot2Token model across a variety of protein-related tasks, achieving better or competitive results. Also, in some tasks, multi-task learning boosts the performance and makes the training stable (Tables 2, 5 and 11). This framework excels in unifying diverse tasks into a general next-token prediction format, which can significantly reduce the cost of training and development compared to specialized models. Moreover, the development of S-ESM, a structure-aware version of the ESM model, highlights Prot2Token’s ability to capture 3D structural information from protein sequences, thereby upgrading the base model to be structure-aware. However, Prot2Token faces challenges in tasks such as 3D structure prediction, particularly in encoding and decoding 3D structural information. While the current method, FoldSeek, effectively encodes 3D structures into discrete tokens, it cannot reverse the process and decode these tokens back into 3D structures.

Currently, Prot2Token connects a PLM to a decoder language model that is initialized using random weights. We believe that integrating a pre-trained PLM into a pre-trained LLM, such as the LLaMA models (Touvron et al., 2023), could further enhance its capabilities in advanced PLP tasks. This approach parallels the development of large vision-language models, like LLaVA (Liu et al., 2024), GPT-4V (Achiam et al., 2023), and Chameleon (Team, 2024), and can lead to more accurate utilization of protein sequence understanding. Additionally, this integration could improve tasks such as protein-protein interface prediction by better understanding the implicit inductive biases in the labels from the limited samples.

## A. Appendix

### A.1 Related Work

We categorize PLP into four categories: sequence-based models, structure-based models, and models that integrate both sequence and structure information, and also, a new category of models named dialogue-based protein language models.

#### Sequence-based model

These models are predominantly sequence-based, yet they are not restricted to this approach alone. Protst employed biomedical textual data (Xu et al., 2023) They used a multi-task learning approach to learn from different types of tasks (unimodal mask prediction, multimodal representation alignment and multimodal mask prediction) simultaneously and then applied their model to different downstream tasks. TAPE (Rao et al., 2019) employed self-supervised pretraining on large protein sequences datasets and fine-tuning it on specific tasks to predict protein properties. Ankh (Elnaggar et al., 2023) utilized protein sequences as input and generates predictions related to protein structure and function. ProGen2 (Madani et al., 2023) generated protein sequences with protein sequences and controllable tags specifying protein properties.

#### Structure-based model

Some papers tried to create PLM models which are more structure-aware. For example, Saprot (Su et al., 2023) used structure-aware vocabulary that combines residue and 3D geometric feature with ESM backbone. Also, (Wang et al., 2022b) developed a structure-aware model with multi-tasking capabilities, using prompts to guide the model’s focus on different structural levels of proteins. GVP (Jing et al., 2020) used 3D protein structures represented as graphs where nodes correspond to amino acids and edges represent spatial proximity. The model was evaluated on two key tasks: computational protein design (CPD), which predicted properties for individual amino acids, and model quality assessment (MQA), which predicted global properties of the protein structure. GearNet (Zhang et al., 2022) used protein structures represented as residue-level relational graphs as input and tried to have fold classification and function prediction. CDConv (Fan et al., 2022) used protein data consisting of 1D sequences and 3D geometric coordinates of amino acids. The model evaluated on four key tasks: protein fold classification, enzyme reaction classification, gene ontology term prediction, and enzyme commission number prediction

#### Combination of sequence and structure

Some other methods merged both sequence and 3D structure information. CoupleNet (Hu et al., 2023) integrated protein sequence and structure information and created a framework for these two types of data, allowing the network to learn complex representations of proteins by leveraging both sequence and structural data. S-PLM (Wang et al., 2024) also used both contact maps for structure information and sequences together and employed a contrastive loss function to transfer information between sequence and 3D structure. And Prostt5 (Heinzinger et al., 2023) proposed a bilingual language model designed for protein sequences and structures. They used language modeling techniques to simultaneously process and translate between one-dimensional amino acid sequences and three-dimensional protein structures. Moreover, Prot2Text (Abdine et al., 2023) focused on predicting a protein’s function by combining the 3D structure using a graph neural network (GNN) and LLM in an encoder-decoder framework, Prot2Text. Their multimodal approach allowed for generating detailed and accurate protein function descriptions in a free-text style as the output of the model for enhancing the understanding of proteins’ functionalities. DeepFRI used protein sequences and 3D structures tried to predict protein functions with providing site-specific annotations at the residue level. they used graph convolutional network that integrates sequence features with structural information. LM-GVP (Wang et al., 2022a) used amino acid and 3D protein structure as input to predict tasks, including fluorescence, protease stability, and functions derived from GO terms. LM-GVP integrates a protein LLM for sequence information with a GNN for structural information, allowing it to leverage the combined data to improve prediction accuracy. ESM-GearNet (Zhang et al., 2023) introduced three fusion strategies for combining sequence and structure representations: serial fusion, parallel fusion, and cross fusion. It utilized ESM-2, combined with structure encoders like GVP, GearNet, and CDConv. The model was evaluated on tasks such as function annotation and enzyme classification by leveraging the combined sequence-structure data.

#### Dialogue-base protein language model

Recent advancements have introduced dialogue-based protein language models, leveraging LLMs to address PLP tasks, including de novo protein design. ProLLaMA (Lv et al., 2024) integrates protein and NLP capabilities, using a two-stage training framework to adapt a general LLM into a protein LLM, excelling in protein sequence generation and property prediction. InstructProtein (Wang et al., 2023) aligns human and protein languages via knowledge instruction, with bidirectional generation capabilities for predicting textual function descriptions from protein sequences and generating protein sequences from natural language prompts. It employs a knowledge graph-based instruction generation framework to construct high-quality instruction datasets. Despite their promising results in protein generation tasks, these dialogue-based models have not been deeply investigated across the full spectrum of PLP tasks. Their effectiveness remains limited due to the lack of a robust tokenization strategy for diverse PLP tasks.

### A.2. Tokenizer

We have three tokenizers for two encoders and one autoregressive decoder.

#### A.2.1 Encoders

##### ESM-2

This model employs a character-level tokenizer specifically designed for amino acid sequences, where each amino acid is represented by a unique token. Additionally, the tokenizer incorporates special tokens like end-of-sequence (EOS), masking, unknown values, padding purposes, and seven other tokens (Lin et al., 2023). Overall, the tokenizer comprises a total of 33 distinct tokens.

##### BARTSmiles

The BARTSmiles model utilizes a unigram tokenizer specifically designed for the SMILES notation of molecular sequences. It is trained on a big corpus of SMILES, ensuring robust coverage of chemical space. The tokenizer incorporates a vocabulary of 1021 unique tokens, which adequately captures individual logical elements, such as atoms and chemical bond symbols from SMILES strings. Additional tokens are reserved for special purposes like end-of-sequence (EOS), beginning-of-sequence (BOS), padding (PAD), and masking, bringing the total vocabulary size to 1025 tokens.

#### A.2.2. Autoregresive decoder

##### Multi-class classification

Multi-class classification involves categorizing instances into one of several classes, making it a foundational approach for various protein-related tasks. Common examples within this domain include localization, protein family classification, enzyme reaction categorization, and fold classification. For these tasks, labels are transformed into discrete tokens. Take TargetP 2.0 localization task, for instance, which features five distinct classes: signal peptide, mitochondrion, chloroplast, thylakoid, and other (Armenteros et al., 2019). During the tokenization phase, these are converted into the respective tokens “sp”, “mt”, “ch”, “th”, and “other”, each symbolizing a unique localization class for protein sequences.

##### Regression

The labels in this category of tasks are continuous data, represented as either floating-point or integer numbers. Stability prediction and protein-ligand affinity prediction are good examples of this type. There are two approaches to tokenizing floating labels: The first involves measuring the range of labels and dividing it into fixed-sized bins, with each number falling into one of these bins. For instance, in a protein task, where target scores range from 0.0 to 10.0, dividing this range into 1.0-sized bins results in 11 distinct bins. However, we opted for the second approach due to a limitation of the binning method: it is quite common for some bins to have very few or even no samples, leading to imbalanced data representation and potential biases in model training. In the second approach, each floating number is encoded into several single digits, offering a more granular and balanced representation of numerical values, ensuring a more uniform distribution of data across the model (Flam-Shepherd & Aspuru-Guzik, 2023). For example, a protein property measured as -0.65 is tokenized into a sequence like {“minus”, “0”, “.” “6”, “5” }, representing the sign, integer part, dot, and fractional digits, respectively. For the training stage, we considered four decimal places for all regression labels.

##### Multi-label classification

In tasks such as Deeploc 2 sub-cellular localization (Thumuluri et al., 2022), EC, and GO, proteins may be classified into multiple categories simultaneously, necessitating a distinct tokenization approach. The GO dataset is a prime example of a multi-label dataset, where proteins are categorized based on their biological processes, cellular components, and molecular functions, often resulting in multiple GO terms being assigned to a single protein. To adeptly manage this complexity, our tokenization strategy is designed to represent multiple labels for a single protein sequence. We tokenize each class associated with a protein as a unique and distinct code, capturing the full spectrum of the annotations. For example, considering GO dataset, labels of a protein with GO terms “GO:0005737” (cytoplasm), “GO:0005829” (cytosol), and “GO:0005654” (nucleoplasm) are tokenized as a sequence of *{*“go:0005737”, “go:0005829”, “go:0005654”*}*.

##### Hierarchical classification

In tasks such as EC and ER predictions, proteins are categorized hierarchically. For EC, each enzyme is assigned a series of numbers representing its specific catalytic activity. If the goal is to do hierarchical classification, it necessitates a specialized tokenization approach. As an example, the EC classification system is divided into four levels: the first level indicates the main enzyme class, the second level specifies the subclass, the third level defines the sub-subclass, and the fourth level denotes the serial number of the enzyme in its sub-subclass. We tokenize each EC number associated with an enzyme into a hierarchical sequence of tokens. For example, an enzyme with EC numbers “1.1.1.1” and “2.2.2.2” is tokenized as {“ec 1”, “1”, “1”, “1”, “ec 2”, “2”, “2”, 2” }, with each part of the EC number being represented as an individual token. This approach allows the model to capture the hierarchical nature of enzyme classifications effectively, ensuring that the different levels of EC labels are properly represented and learned. In addition to this hierarchical tokenization, we could employ a second approach where each complete EC number is treated as a unique and distinct code similar to GO datasets. For example, an enzyme with EC numbers “1.1.1.1” and “2.2.2.2” could be tokenized as {“ec 1 1 1 1”, “ec 2 2 2 2”} , with each token acting as a representative for an entire EC number. This method is also applicable to the ER dataset. This alternative tokenization could yield different results depending on the task. In our early experiments, we found that converting ER labels into a hierarchical format reduced performance compared to using a multi-label classification format, while the opposite was true for the EC task. However, we did not investigate this thoroughly in our work.

The labels of certain tasks in our model are directly correlated with the number of amino acids present in the input protein. Tasks such as 3D structure, post-translational modification (PTM), and secondary structure (SS) prediction are prime examples of this correlation. In the sections that follow, we detail our approach to tokenizing these tasks, highlighting how the specific characteristics of each protein, reflected in its amino acid sequence, inform the respective tokenization processes.

##### 3D structure

In this task, the 3D structure of proteins is converted into discrete 3Di tokens using the FoldSeek (van Kempen et al., 2023) method. Specifically, this method transforms each protein structure into 20 types of tokens, corresponding to each amino acid it contains. For instance, if a protein is composed of 100 amino acids, the FoldSeek method translates its 3D structure into 100 3Di tokens that are aware of the structural configuration. This approach not only preserves the essential spatial information of the protein’s 3D structure but also facilitates the tokenization process by providing a structured and interpretable representation of complex molecular shapes.

##### Secondary Structure

The approach to tokenizing SS prediction in proteins is akin to the method used for 3D structures. In this task, each amino acid is converted into a token representing its secondary structure, categorized as either alpha helix, beta strand, or random coil. For instance, consider a peptide sequence ACDEFGHIKLMNPQRSTVWY. The corresponding secondary structure might be represented as HHHHHHCCCEEEEEEECCC, where each letter corresponds to a specific structural form – “H” for alpha helix, “C” for random coil, and “E” for beta strand.

##### Post Translational Modification

In the realm of PTM, the tokenization process is intricately linked to the protein’s amino acid sequence, with each amino acid evaluated for potential modifications, including phosphorylation, methylation, and acetylation, among others. To tokenize PTM labels, our methodology involves identifying and indexing all potential modification sites specific to each PTM type. For instance, in tasks focusing on phosphorylation, amino acids like serine (S), threonine (T), and tyrosine (Y) are potential phosphorylation sites. We represent these sites through a series of tokens that differentiate between all potential and positive sites. This is achieved by delineating the indices of these sites in a list, separated by a special token, *<sep >*, to distinctly mark the transition. Consider a protein sequence “ASSKYKAMTV”; the target tokenization for phosphorylation might be represented as {“2”, “3”, “5”, “9”, “*<sep >*“, “3”, “9”} , where the numbers before *<sep >* indicate the potential sites, and those after *<sep >* denote the actual modification sites (Figure 4).

**Figure 4.**
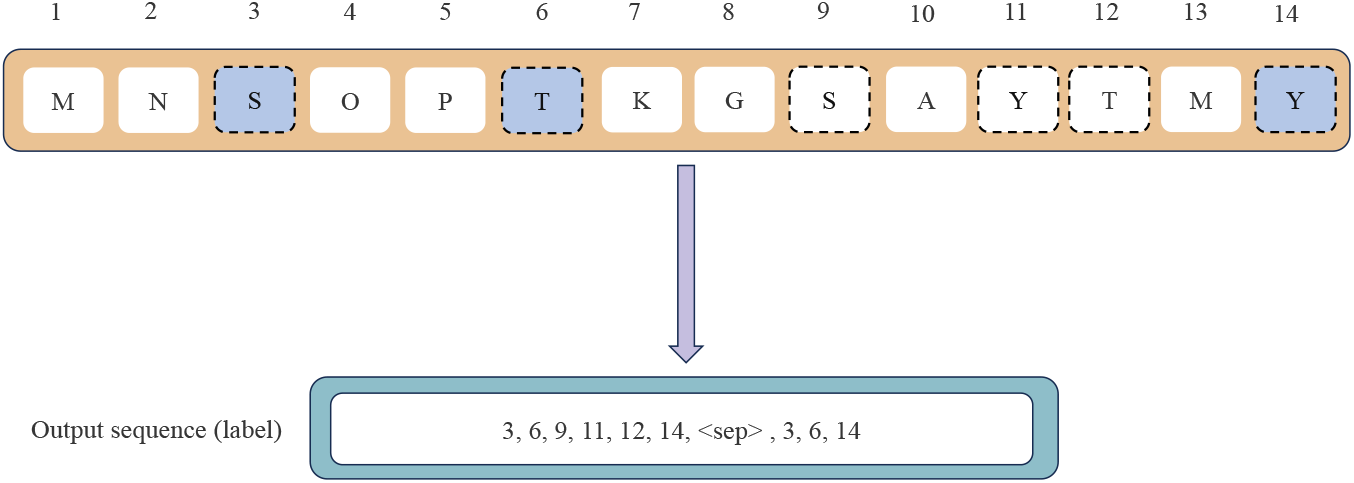
Illustration of PTM tokenization for phosphorylation. The protein sequence is shown with all potential phosphorylation sites (Serine, Threonine, and Tyrosine) highlighted. The output sequence (label) represents the indices of all potential sites, followed by the actual modification sites, separated by a special token *<*sep*>*.

##### Protein-Protein Interface

The tokenization process for the protein-protein interface task prediction involves converting the 3D structural information of protein complexes into a format suitable for sequence-based models. As illustrated in (Figure 5), the process begins with the 3D structure of a protein complex. This structure is then represented as a binary interaction matrix, where each row and column corresponds to specific amino acids from the interacting proteins. A cell in the matrix contains a 1 if there is an interaction between the corresponding amino acids and a 0 otherwise. To use autoregressive modeling, the interaction matrix is transformed into a sequence of coordinate pairs. Each pair (i, j) in the sequence denotes the indices of interacting amino acids from the two proteins. This sequence of pairs effectively captures the interaction information in a tokenized format that can be processed by the Prot2Token framework.

**Figure 5.**
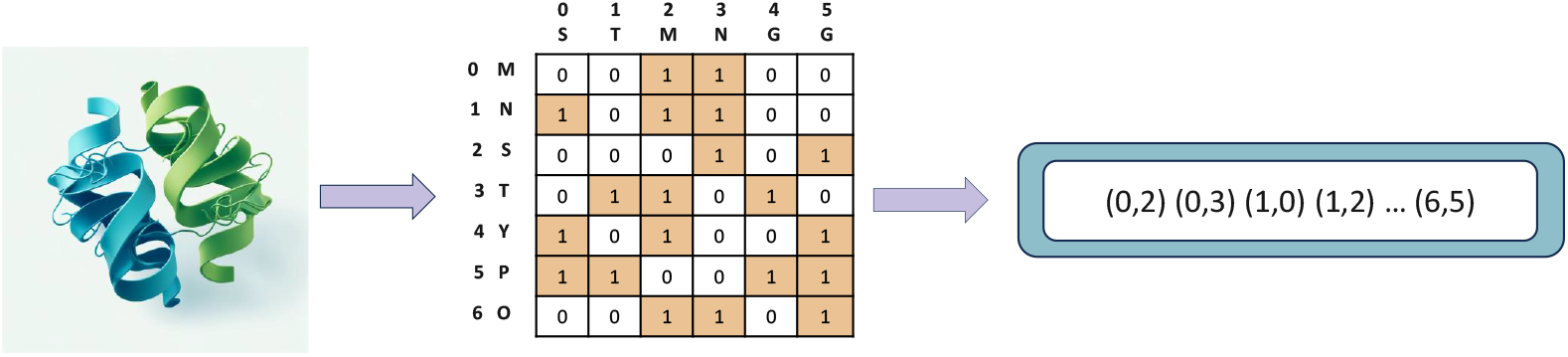
The image illustrates the process of tokenizing protein-protein interface labels. It starts with a 3D structure of a protein complex on the left. This 3D structure is converted into a binary interaction matrix, shown in the center, where rows and columns represent different amino acids from the interacting proteins. Each cell in the matrix indicates whether there is an interaction between the corresponding amino acids (1 for interaction, 0 for no interaction). The matrix is then transformed into a sequence of coordinate pairs on the right, where each pair (i, j) denotes the indices of interacting amino acids from the two proteins.

### A.3. Architecture

The Prot2Token framework integrates multiple components to handle various protein-related tasks within a unified architecture. It includes two primary encoders: the chemical encoder and the protein encoder, each equipped with a bidirectional attention prefix to process their respective inputs. The chemical encoder processes chemical sequences, i.e., SMILES representations of molecules, while the protein encoder handles protein sequences, both converting these inputs into feature embeddings. Each input pair of protein and SMILES sequences is associated with a specific task token that guides the model on the specific task it is addressing. The encoded features from both the chemical and protein encoders are concatenated together and then pass through a linear layer to reduce the size, followed by cross-attention layers to connect to the decoder, an autoregressive language model. The decoder receives the concatenated features from both encoders along with the task tokens to generate the target sequences, one token every time, sequentially. The input to the decoder includes a special beginning-of-sequence (BOS) token followed by the task token. The predicted sequence is then compared to the target sequence for loss calculation, which guides the training process. This architecture allows Prot2Token to effectively unify various tasks into a next-token prediction framework, leveraging multi-task learning across different PLP tasks. We build two Prot2Token models based on the configuration in Table 7.

**Table 7.** Prot2Token model configurations.

| ARCHITECTURE | ENCODER | DECODER |  |  |  |
| --- | --- | --- | --- | --- | --- |
|  |  | EMBEDDING DIMENSION | FEEDFORWARD DIMENSION | HEADS | LAYERS |
| PROT2TOKEN (ESM-35M) | ESM-35M | 480 | 960 | 8 | 4 |
| PROT2TOKEN (ESM-650M) | ESM-650M | 640 | 1280 | 8 | 4 |

#### Autoregressive language model

Autoregressive language modeling is a computational approach mostly used in NLP that predicts subsequent tokens in a sequence based on the preceding tokens. This method operates on the principle of conditional probability, wherein each token is generated one after another, with the prediction of each new token being influenced by the sequence of tokens that came before it. Autoregressive models, such as GPT (Radford et al., 2018), learn these probabilities by being trained on vast datasets of text, allowing them to generate coherent and contextually relevant text sequences. This approach is distinct for its sequential nature, contrasting with autoencoding models that predict missing tokens in a sequence.

We used task prompts to handle multiple tasks during one training session. This unique prompt token serves as a clear indicator of the model, specifying the type of task that the decoder needs to address, by learning its embedding during the training process (Figure 6B). The role of the task token is crucial; it guides the decoder to adjust its sequence predictions to fit the specific requirements of the task at hand. Interestingly, while the decoder is informed by these task tokens as a prompt, the encoders operate without explicit knowledge of the target task based only on the protein and chemical sequences. This approach implicitly functions as a regularizer during joint training, enhancing the model’s performance. We define “joint training” in a manner akin to what is depicted in Figure 6B, where multiple tasks are combined and trained simultaneously, utilizing task tokens to distinguish and manage each specific task. The benefit of this strategy compared to classical multi-task learning is that we can merge multiple datasets of various tasks at once without having labels of all tasks for every protein. When we do not want to use joint training, we can remove the task token similar to Figure 6A.

**Figure 6.**
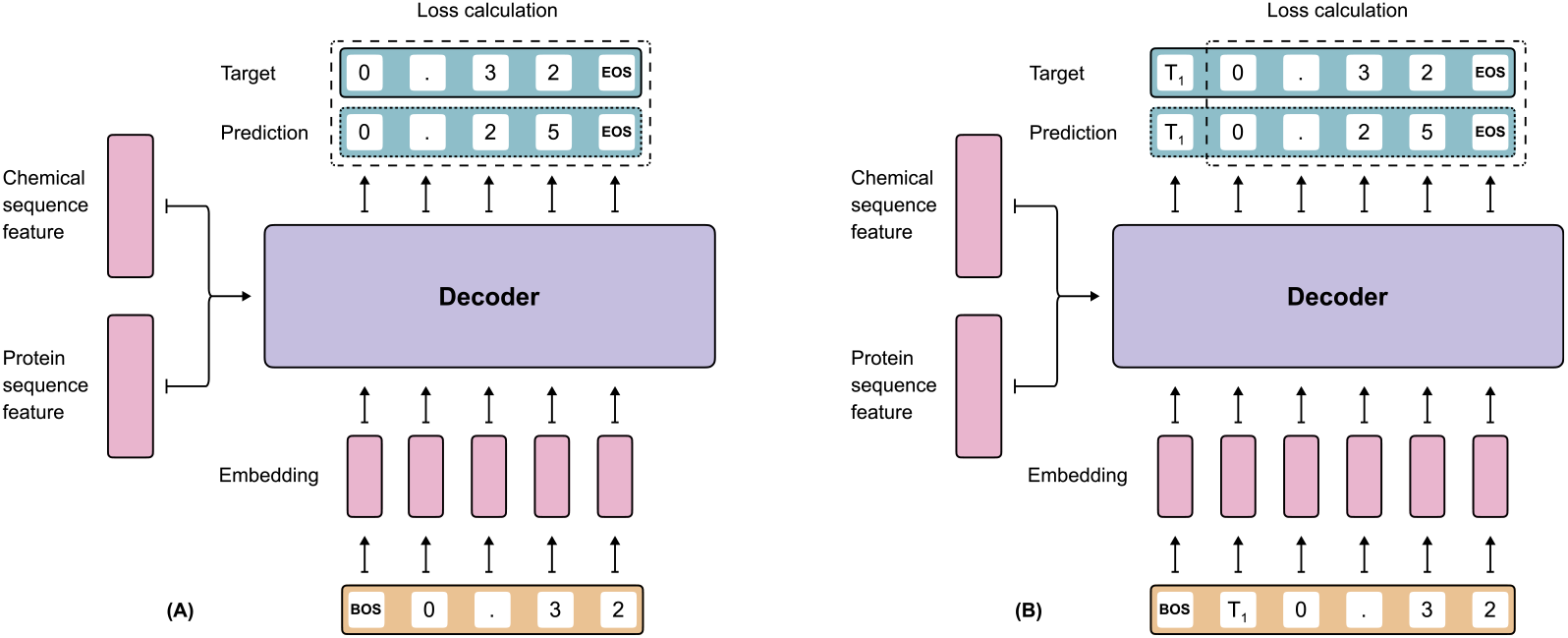
Training dynamics of the Prot2Token decoder. During the training phase, the decoder receives features from the protein and chemical encoders via cross-attention and uses the embedded label tokens as input to predict the subsequent tokens. (A) demonstrates the training process when Prot2Token is applied to a single task, where the task token is omitted, aligning the training methodology with that of a traditional autoregressive language model. (B) illustrates the training setup for multiple tasks, utilizing a unique task token, e.g., T1, as the prompt for each sample of the decoder input, but it is excluded from the loss calculation.

This approach is formulated as follows: Let *T* be the task token, and (*y*_1_, *y*_2_, … , *y*_*N*_ ) be the sequence of target tokens. The probability of the sequence given the task is modeled as *P* (*y*_1_, *y*_2_, … , *y*_*N*_ |*T* ). However, during the calculation, the task token *T* is assigned a weight of zero, effectively excluding it from influencing the probability computations directly. This can be mathematically represented as Equation (1).

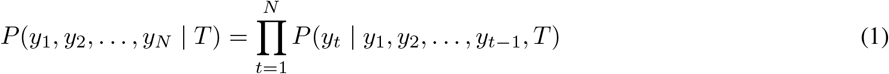

with the weight of *T* in the computation being zero. This ensures that while the task token guides the overall direction of the sequence generation, it does not artificially skew the probabilities of the protein’s label tokens. The primary training objective of the Prot2Token model is to maximize the predictive accuracy of sequences of labels for various tasks while effectively integrating the task-specific guidance provided by the prompt (task) token. This objective is achieved through a carefully designed training process that balances the model’s adaptability to different tasks with its ability to accurately predict sequences of labels. The model is trained to maximize the likelihood of the correct label tokens given a task token. Mathematically, this is represented as maximizing the conditional probability of the sequence given the task token, *P* (*y*_1_, *y*_2_, … , *y*_*N*_|*T* ), where *T* is the task token and (*y*_1_, *y*_2_, … , *y*_*N*_ ) are the label tokens. Formally, the objective is maximized and presented in Equation (2).

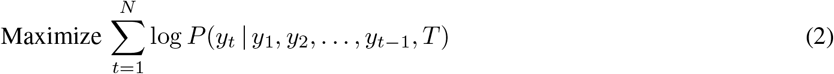

Where the influence of the task token *T* in the computation is acknowledged but its loss weight is set to zero during the calculation of the loss function.

For single-task training, the probability of the sequence is formulated as the product of the conditional probabilities of each label token in the sequence without the influence of a task token. This is mathematically represented as Equation (3).

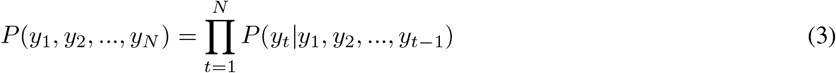

And the optimization objective during training is to maximize the sum of the log probabilities of each label token given the previous label tokens in the sequence. This is expressed as Equation (4).

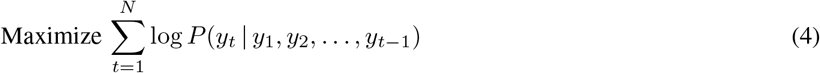

### A.4. Dataset

#### PEER

It presents the PEER benchmark (Xu et al., 2022), a comprehensive and multi-task benchmark for protein sequence understanding. It encompasses diverse tasks such as protein function prediction, localization prediction, structure prediction, PPI, and Protein-Ligand Interaction prediction. The benchmark utilizes various datasets for each task category, ensuring a wide coverage of biological aspects. It is designed to evaluate the performance of different sequence-based methods including traditional feature engineering, various sequence encoding methods, and large-scale pre-trained protein language models. In this work, we consider human PPI, secondary structure, fluorescence, stability prediction and Protein-Ligand Affinity (PLA) datasets from PEER.

#### ProteinShake

The paper (Kucera et al., 2023) introduces ProteinShake, a Python package designed for the creation and evaluation of datasets in deep learning for protein structures. It allows users to easily generate custom datasets or use pre-processed ones from the Protein Data Bank (PDB) and AlphaFoldDB. Each dataset is associated with prediction tasks and evaluation functions, covering a broad spectrum of biological challenges. ProteinShake also offers standardized data splits based on sequence and structure similarity, and a benchmark demonstrating the impact of pre-training and different data modalities (graphs, voxel grids, or point clouds) on model performance. The tool simplifies accessing protein structure data and standardizes model comparisons, providing a platform for challenging benchmark settings with real-world implications. We consider Protein Family and Structure Similarity datasets from ProteinShake. We use the “structure split” strategy with a similarity threshold of 70% for the evaluation.

#### AlphaFoldDB

AlphaFold Protein Structure Database significantly expands structural coverage in protein-sequence space. It utilizes AlphaFold’s AI-powered predictions to offer a comprehensive database of high-accuracy protein structures. The initial release features over 360,000 predicted structures covering 21 model-organism proteomes and at the time of writing, it expands over 200 million proteins. AlphaFold DB is notable for its extensive coverage, including most sequences from the UniRef90 dataset, and provides a valuable resource for researchers in various biological and biomedical fields. For our work, we consider the prediction of 542,378 proteins of the SwissProt (Bairoch & Apweiler, 2000) database.

#### CATH dataset

The preprocessed protein sequences with CATH annotations were downloaded from the literature (Wang et al., 2024). Specifically, the CATH nonredundent S40 (release v4 3 0) is used, whose proteins of maximally 40% sequence similarity and only one represented sequence with the longest sequence length was selected from one CATH superfamily.

#### Phosphorylation

The phosphorylation dataset is downloaded from (Wang et al., 2017) and modified on serine (S) and threonine (T) amino acids. The dataset has been annotated by UniProt/Swiss-Prot and used as positive data, while the same amino acid excluding annotated phosphorylation sites from the same proteins were regarded as negative data. The testing set has no more than 50% similarity with the training and validation set.

#### Auxiliary self-supervised tasks

We also create auxiliary self-supervised tasks. In these auxiliary tasks, we supplied sequences of amino acids, with the objective being to pinpoint the positions of specific amino acid types. For example, in sequences containing the amino acid ‘S’, such as “ASGTSMYK”, we would label the locations of ‘S’ as the target, resulting in a sequence of indices like {2, 5} . At the end, we craft 20 auxiliary self-supervised tasks given each amino acid as one task. The important point about these types of tasks is that as long as we have access to protein sequences, they are free to craft, and therefore, no human labeling is required.

Other than the mentioned datasets, we use GO (Consortium, 2008), ER (Webb et al., 1992), EC (Omelchenko et al., 2010), Fold classification (Hou et al., 2018), Target-P 2.0 localization (Armenteros et al., 2019) datasets. The localization has 13,005 samples from 5 different categories as well as their cleavage site positions. The statistics of other datasets that we use are placed in table Table 8.

**Table 8.**
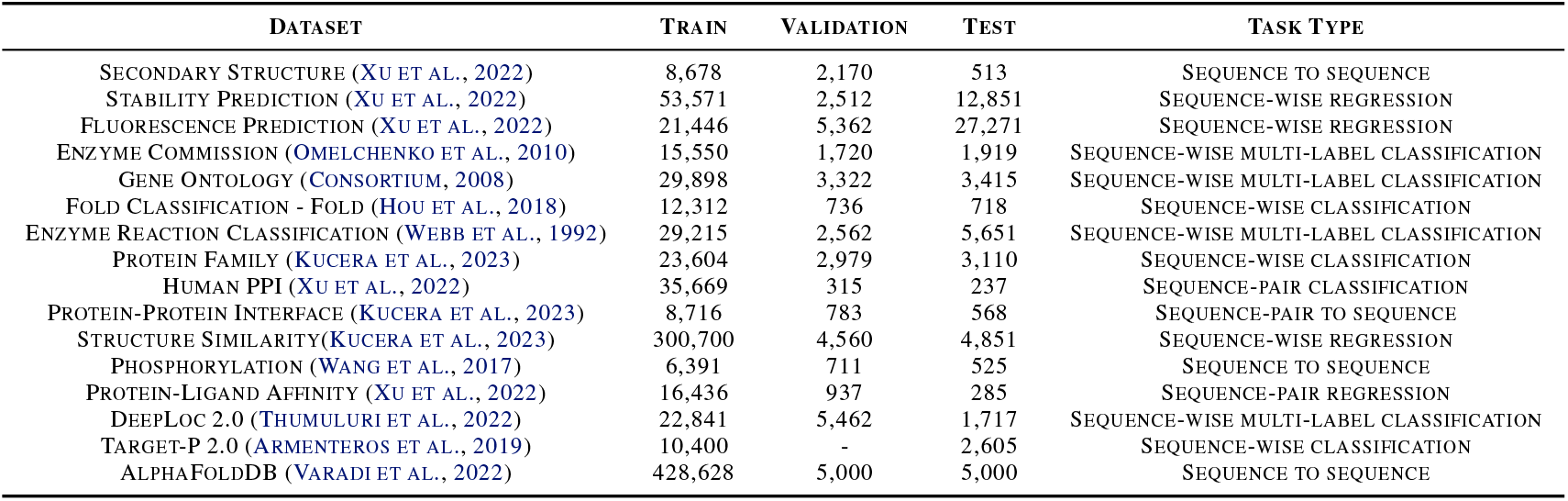
Dataset Statistics Overview. This table presents the details of the datasets utilized in this study. We employ a structural split approach with a maximum of 70% similarity, for protein family classifications, protein-protein interface and structural similarities datasets from ProteinShake.

#### A.4.1 Prepare S-ESM dataset

In our study, we utilized the AlphaFold database (Jumper et al., 2021; Varadi et al., 2022), which comprises 542,378 Swiss-Prot proteins. Initially, we excluded proteins with an average pLDDT score below 80% and those exceeding 1022 amino acids. This filtering process resulted in a refined dataset of 438,628 proteins. Further, we focused on proteins with an average pLDDT score above 90%. From this subset, we segregated 5,000 proteins each for the validation and test sets. The remaining 428,628 proteins were designated for the training set. Furthermore, we converted all 3D structures of training, validation, and test sets to 3D aware sequences using the FoldSeek method (van Kempen et al., 2023) and considered them as the target labels.

### A.5. Additional Experiments

#### A.5.1. Phosphorylation site prediction

In our initial attempts with the PTM-Phosphorylation task (Esmaili et al., 2023) using the Prot2Token model, we focused on predicting positive phosphorylation sites but found the performance unsatisfactory. Initially, we attributed this to the label structure, leading us to modify the label format as described in the methods section. This adjustment yielded a slight improvement in our metrics, yet the model’s performance remained suboptimal. We then considered that the issue might stem from the lack of inductive biases in the Prot2Token model, biases that specialized model’s inherently possess. Our baseline approach was akin to a named entity recognition (NER) task, where a feedforward layer was added to the model to classify potential phosphorylation sites among amino acids. This method essentially narrowed the problem’s search space in two ways: firstly, by classifying amino acids into categories using softmax, and secondly, by limiting the classification to potential phosphorylation sites such as the amino acids S and T. Recognizing that the Prot2Token model does not intrinsically include these biases, we decided to integrate a set of simple self-supervised auxiliary tasks into the main training process, to help the model learn these biases in its prediction effectively.

Our empirical data in Table 9 suggests a direct correlation between the number of auxiliary samples and the improvement in phosphorylation task performance. Notably, expanding the scope of auxiliary tasks to include amino acids KNR, in addition to STY, as the self-supervised tasks marked the most significant performance enhancement. Given that generating auxiliary samples from raw protein sequences is a cost-free process, it is worthwhile to investigate the extent to which this strategy can further enhance performance.

**Table 9.**
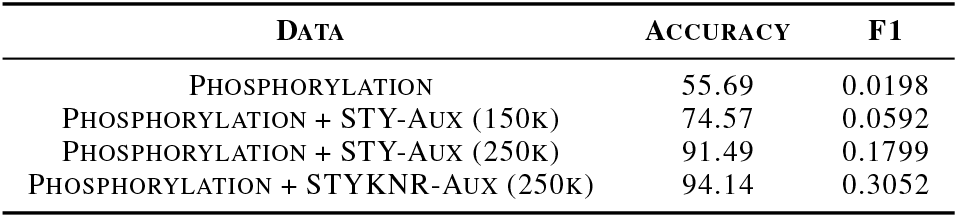
Phosphorylation prediction with Prot2Token. “Aux” denotes self-supervised auxiliary tasks. All results are based on Prot2Token (esm-650m) model.

#### A.5.2. Stability Prediction and Fluorescence

These tasks focus on determining the stability and fluorescence properties of a protein within a specific environment. We selected the PEER database, which includes distinct training, validation, and test sets for those tasks. In our comparative analysis, we maintained the ESM model weights as fixed and only unlocked the last six layers of it to be fine-tuned and connected it to the decoder. Also, for the baseline, we consider a linear regression layer as its head. In addition, given that the Spearman correlation metric is insensitive to normalization, we normalized the labels to fall within a 0 to 1 range and compared it with unnormalized, there was a significant improvement. It indicates that the decoder part of the model needs to learn the structure of regression output at first to have a better prediction and by doing normalizing, the model learns the structure of output faster.

#### A.5.3. Protein-Ligand Affinity

This task is similar to the stability and fluorescence tasks in terms of output, with the main difference being the input: each sample includes a protein sequence and a SMILES representation of a molecule. For the protein encoder, we kept the ESM model weights fixed, unlocking only the last six layers for fine-tuning. In contrast, for the chemical decoder, we found that fine-tuning all layers of BARTSmiles, except the embedding weights, yielded the best performance. We used the PEER database for this task, which provides distinct training, validation, and test sets. Additionally, as with the other regression tasks, the labels were normalized.

#### A.5.4. TargetP AND Cleavage site

We utilized the TargetP-2 dataset for our localization studies, which encompasses both cleavage site data and five types of localization labels. We represented the label format as a combination of classification and regression tasks, for instance, {“sp”, “96” }, where “sp” denotes the localization label (Signal Peptide) and “96” indicates the cleavage site’s location. Additionally, to evaluate the model, we implemented a 5-fold cross-validation strategy. We considered fine-tuning only the last layer of the ESM models for both the Prot2Token model and the baseline comparison. Table 10 presents a comparative analysis of Prot2Token against ESM with a linear classifier head. The results suggest that by enabling the model to learn the locations of different amino acids through self-supervised auxiliary tasks, it achieves more accurate predictions of cleavage site positions. Furthermore, the performance in localization prediction also shows improvement with the integration of auxiliary tasks. We attribute this enhancement in performance to the model’s improved understanding of cleavage site positions. Note that the performance of bigger models was very similar to the smaller ones.

**Table 10.** Localization and cleavage site prediction. “Aux” denotes self-supervised auxiliary tasks using STYKNR amino acids. Localization and cleavage site metrics are based on Macro-F1 and MAE, respectively.

| METHOD | AUX-TASKS | CLEAVAGE SITE | LOCALIZATION | MODEL |
| --- | --- | --- | --- | --- |
| ESM-2 | - | - | 90.96 | ESM-35M |
| OUR | - | 3.6392 | 90.56 | PROT2TOKEN (ESM-35M) |
| OUR | AUX-STYKNR (12K) | 2.9205 | 92.30 | PROT2TOKEN (ESM-35M) |

**Table 11.**
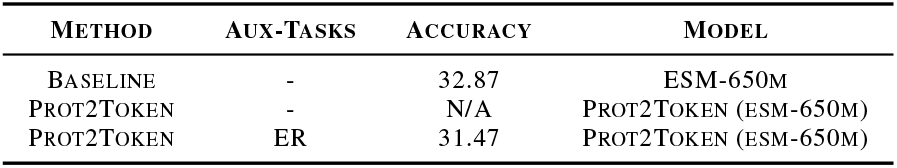
Fold classification training in single-task and multi-task training on Fold-fold test set.

#### A.5.5. Fold classification

For this task, we maintained the ESM model weights as fixed and only unlocked its last six layers of it to be fine-tuned and connected to the decoder. Many classes in this dataset have a low number of samples, e.g., one sample for a high number of classes. That is why we saw unstable training when we did single-task training on Prot2Token. However, when we combined Fold classification with auxiliary tasks like ER, the training became stable (Table 11).

#### A.5.6. Human Protein-Protein Interaction

For this task, we maintained the ESM model weights as fixed and only unlocked the last four layers of it to be fine-tuned and connected to the decoder. Note that to give the encoder two sequences at one feed for PPI, we concatenated two sequences using the EOS token. We observed that adding more tasks helped boost the performance of Human PPI (Table 12). However, Prot2Token tended to overfit on this task, indicating that the improvement from adding auxiliary tasks may be due to the regularization effect of multi-task learning.

**Table 12.** Human PPI performance on PEER test set.

| METHOD | AUX-TASKS | ACCURACY | MODEL |
| --- | --- | --- | --- |
| PEER (FINE-TUNED) | - | 78.17 | ESM-1B |
| PROT2TOKEN | - | 71.3 | PROT2TOKEN (ESM-650M) |
| PROT2TOKEN | DEEPLOC | 78.48 | PROT2TOKEN (ESM-650M) |
| PROT2TOKEN | DEEPLOC+ER+FOLD | 80.17 | PROT2TOKEN (ESM-650M) |

#### A.5.8. Gene Ontology and Enzyme Commission

For the GO and EC tasks, we encountered a limitation in calculating the Fmax metric, which is commonly used for performance evaluation in these tasks. Instead, we used accuracy and F1 score to assess our model’s performance. Consequently, we were unable to directly compare our results with those of other methods that report their performance in terms of Fmax. This discrepancy highlights a significant challenge in benchmarking our approach against existing methods. The GO tasks are further divided into three categories: biological process (BP), molecular function (MF), and cellular component (CC). We jointly trained all four tasks (the three GO tasks and the EC task) together in a multi-task learning manner. Detailed performance metrics for these tasks are presented in Table 13. We maintained the ESM model weights as fixed and only unlocked the last four layers of it to be fine-tuned and connected it to the decoder and a linear classifier for Prot2Token. Note that labels in these tasks are highly imbalanced.

**Table 13.**
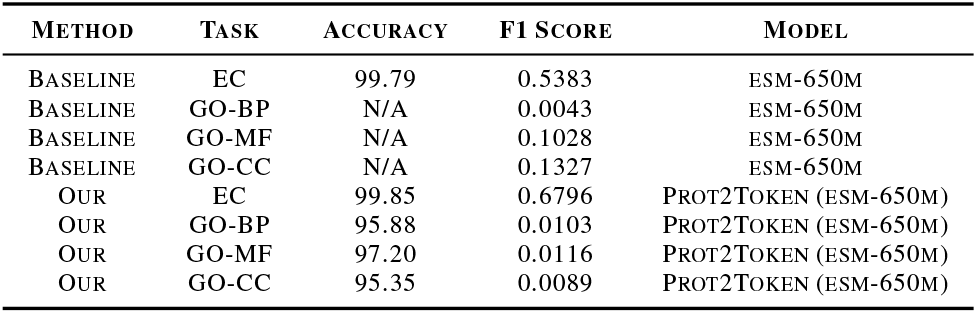
Comparing GO and EC tasks with the baseline on accuracy and F1 score metrics. The baseline is a linear evaluation of ESM.

**Table 14.** S-ESM on multiple structure-related PLP tasks. In both ESM and S-ESM models, we appended a linear classifier to the backbone for classification.

| TASK | ESM (LE) | S-ESM (LE) | S-ESM (FT) | COUPLENET<br>(HU ET AL., 2023) | GEARNET<br>(ZHANG ET AL., 2022) | PROTEINSHAKE<br>(KUCERA ET AL., 2023) |
| --- | --- | --- | --- | --- | --- | --- |
| GO-BP (FMAX) | 0.339 | 0.4139 | **0.4834** | 0.467 | 0.356 | - |
| GO-CC (FMAX) | 0.3976 | 0.4746 | 0.4473 | **0.494** | 0.414 | - |
| GO-MF (FMAX) | 0.434 | 0.5371 | **0.6858** | 0.669 | 0.503 | - |
| EC (FMAX) | 0.8002 | 0.8111 | **0.8721** | 0.866 | 0.73 | - |
| FOLD-FAMILY (ACC) | 98.98 | 98.74 | 98.66 | **99.7** | 95.3 | - |
| FOLD-SUPER FAMILY (ACC) | 67.38 | 74.08 | 76.32 | **82.1** | 42.6 | - |
| FOLD-FOLD (ACC) | 32.87 | 39.14 | 40.39 | **60.4** | 28.4 | - |
| PROTEIN FAMILY (ACC) | 69.39 | **73.76** | 71.83 | - | - | 41.4 |
| STRUCTURE SIMILARITY (SPEARMAN) | 0.4027 | 0.3984 | 0.4743 | - | - | **0.573** |
| ER (ACC) | 83.81 | 84.09 | 85.17 | **89.0** | 79.4 | - |

#### A.5.8. Protein-Protein Interface

The performance of the Prot2Token model on the protein-protein Interface task was not satisfactory. In the first attempt, the model struggled to learn the structure of the labels for this task. To address this issue, we added auxiliary self-supervised tasks to support the learning process, but this did not result in significant improvement. We believe that this low performance is primarily due to the lack of inductive biases in the decoder for understanding the structure of the output labels, exacerbated by the low number of samples available for this task. Using a pre-trained language model instead of the current randomly initialized one could potentially solve this lack of understanding problem, presenting a good direction for future research to examine the benefits and limitations of this approach.

### A.6. Structure-Aware ESM

FoldSeek (van Kempen et al., 2023) effectively compresses 3D structures into 3Di tokens, facilitating rapid and precise protein structure searches. However, a notable limitation of FoldSeek is the challenge of reversing the tokenization process, which hinders its direct application in predicting protein 3D structures. We prepared the training dataset as described in Appendices A.2.2 and A.4.1. We fine-tuned the last 22 layers of the ESM-650m model connected to the decoder. After completing 16 epochs of training (Figure 7), we assessed the fine-tuned encoder’s representational capabilities using the CATH dataset, inspired by S-PLM (Wang et al., 2024). This evaluation aimed to demonstrate that the sequence embeddings generated by Prot2Token are aware of structural information and can effectively distinguish between different structure domains.

**Figure 7.**
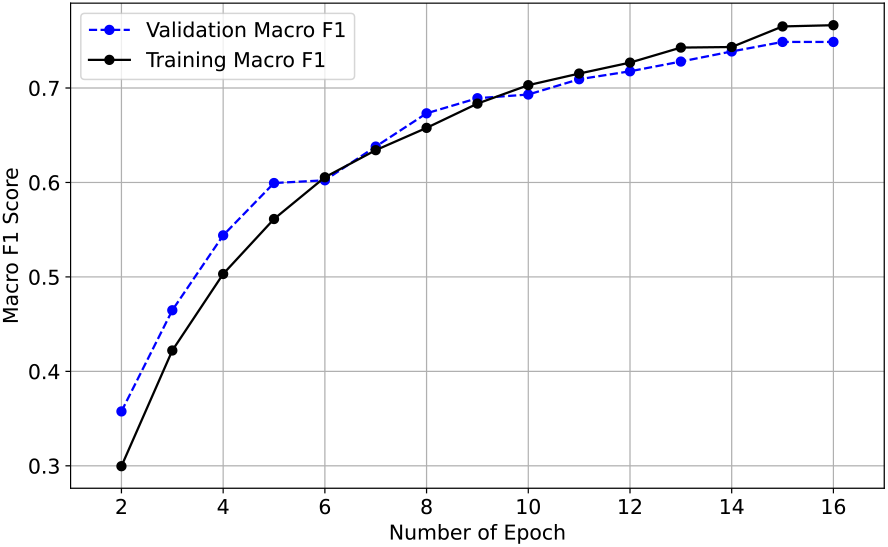
Training process of the Prot2Token model on 3Di tokens derived from the AlphaFold DB. The graph highlights the model’s learning trajectory, as evidenced by the continual increase in the macro F1 score on the validation set. Also, even after 16 epochs of training, there was no sign of overfiting which suggests that we can improve the performance by training further.

We employed T-SNE to visualize the protein sequence embeddings generated by the last layer of both ESM and S-ESM, reducing the original 1280D sequence embeddings to 2D embeddings. In the main body of the paper, Figure 3 shows the 2D visualization using the CATH protein sequences. The representations generated by ESM are intertwined for the alpha and beta proteins, whereas those generated by S-ESM are separated based on structural classes. This observation indicates that the embeddings of protein sequences generated by S-ESM are aware of structural information.

Furthermore, we conducted k-means clustering on the embedding of sequence and calculated the adjusted Rand index (ARI) by comparing the predicted clusters with the known CATH classes in the 2D reduced dimension. The ARI calculated using ESM embeddings is -0.002, while the ARI for S-ESM is 0.144. This demonstrates S-ESM’s superior performance in separating CATH structure domains compared to the original ESM weights.

